# A Fast and Accessible Method for the Isolation of RNA, DNA, and Protein to Facilitate the Detection of SARS-CoV-2

**DOI:** 10.1101/2020.06.29.178384

**Authors:** Jose Carlos Ponce-Rojas, Michael S. Costello, Duncan A. Proctor, Kenneth S. Kosik, Maxwell Z. Wilson, Carolina Arias, Diego Acosta-Alvear

**Affiliations:** Department of Molecular, Cellular, and Developmental Biology, University of California Santa Barbara, Santa Barbara, CA; Neuroscience Research Institute, University of California Santa Barbara, Santa Barbara, CA; Center for BioEngineering, University of California Santa Barbara, Santa Barbara, CA

**Author notes:** These authors contributed equally to this work (Jose Carlos Ponce-Rojas, Michael S. Costello). These authors are co-senior authors (Carolina Arias, Diego Acosta-Alvear).

## Abstract

Management of the COVID-19 pandemic requires widespread SARS-CoV-2 testing. A main limitation for widespread SARS-CoV-2 testing is the global shortage of essential supplies, among these, RNA extraction kits. The need for commercial RNA extraction kits places a bottleneck on tests that detect SARS-CoV-2 genetic material, including PCR-based reference tests. Here we propose an alternative method we call PEARL (<u>P</u>recipitation <u>E</u>nhanced <u>A</u>nalyte <u>R</u>etrieva<u>L</u>) that addresses this limitation. PEARL uses a lysis solution that disrupts cell membranes and viral envelopes while simultaneously providing conditions suitable for alcohol-based precipitation of RNA, DNA, and proteins. PEARL is a fast, low-cost, and simple method that uses common laboratory reagents and offers comparable performance to commercial RNA extraction kits. PEARL offers an alternative method to isolate host and pathogen nucleic acids and proteins to streamline the detection of DNA and RNA viruses, including SARS-CoV-2.

## Introduction

The COVID-19 pandemic has had a devastating social and economic impact worldwide. As the disease continues to spread, global SARS-CoV-2 testing is more urgent than ever. The most widely used reference test for SARS-CoV-2 detection relies on the isolation of viral genetic material followed by PCR-based amplification^1–2^. The first step in this approach is the extraction of viral RNA from human samples. Commercial solid-phase RNA extraction kits that isolate viral RNA are the starting point for PCR-based SARS-CoV-2 reference tests^3^. These kits use silica-based columns to purify viral RNA after disruption of cells and viral particles with proprietary reagents. The global demand for these kits has made them a limiting resource for SARS-CoV-2 testing, fueling the development of alternative SARS-CoV-2 RNA isolation methods and protocols. These alternative approaches include organic solvent-based RNA extraction, and the use of chaotropic agents and proprietary buffer formulations.

TRIzol, a phenol- and guanidine-based reagent routinely used for isolation of RNA, DNA, and protein, has been used to isolate SARS-CoV-2 RNA^4–6^. However, TRIzol extraction is labor intensive, which challenges scaling-up to meet testing demands. Moreover, it requires special considerations for the disposal of organic solvents. A 5-minute RNA preparation method has been recently reported, but it depends on expensive proprietary lysis solutions originally developed for genomic DNA isolation^7^. Recently, direct detection of SARS-CoV-2 in nasopharyngeal swab samples without RNA extraction was reported, indicating that the initial RNA isolation step could be omitted^8–10^. Despite encouraging results, this approach results in reduced sensitivity of downstream quantitative PCR (qPCR)-based detection. On average, this method required an additional 5-7 PCR cycles to reach the detection threshold when compared to reactions templated on purified RNA. Because detection of low viral loads is critical for minimizing false negative results, it is essential that new approaches do not compromise sensitivity. In a more recent report, guanidium chloride was used for sample lysis in nasal swabs obtained from COVID-19 positive patients^11^. Total RNA was subsequently precipitated with isopropanol. This approach conveniently concentrates the RNA, which can increase detection sensitivity in downstream analyses. However, the use of the toxic chaotropic agent guanidium chloride requires special disposal guidelines.

To address the aforementioned shortcomings, we developed a simple technique to isolate nucleic acids and proteins from cells and viruses we call PEARL (<u>P</u>recipitation <u>E</u>nhanced <u>A</u>nalyte <u>R</u>etrieva<u>L</u>). PEARL is fast, easy to perform, and uses common laboratory reagents. Moreover, PEARL allows the downstream detection of specific SARS-CoV-2 viral sequences with comparable sensitivity to that afforded by commercial RNA extraction kits. PEARL can be coupled to nucleic acid amplification or immunodetection methods to detect host and viral RNA, DNA, and proteins from multiple sources. PEARL does not require specialized equipment or highly trained personnel, and offers a low-cost straightforward alternative to facilitate virus detection.

## Results

We designed PEARL to provide a low-cost, column-free approach for the isolation of nucleic acids and proteins that uses common laboratory reagents (Fig. 1A, Supplemental Table 1). PEARL uses a non-ionic detergent-based lysis solution (see Materials and Methods) to disrupt cell membranes and viral envelopes, while simultaneously providing conditions suitable for alcohol-based precipitation and recovery of RNA, DNA, and proteins. To benchmark our method, we extracted RNA from de-identified SARS-CoV-2 positive samples using PEARL or a dedicated RNA extraction kit (QIAamp Mini Elute Virus Spin Kit, Qiagen). Next, we used the isolated RNA to examine the levels of the SARS-CoV-2 nucleoprotein (N) gene as well as the host RNaseP mRNA in the samples using the 1-step reverse transcription qPCR reference test for COVID-19 recommended by the United States Centers for Disease Control and Prevention (CDC) (TaqMan RNA-to-Ct 1-Step Kit, ThermoFisher). In these experiments, we detected the SARS-CoV-2 N1 site using the qPCR primers and probes recommended by the CDC.

**Figure 1.**
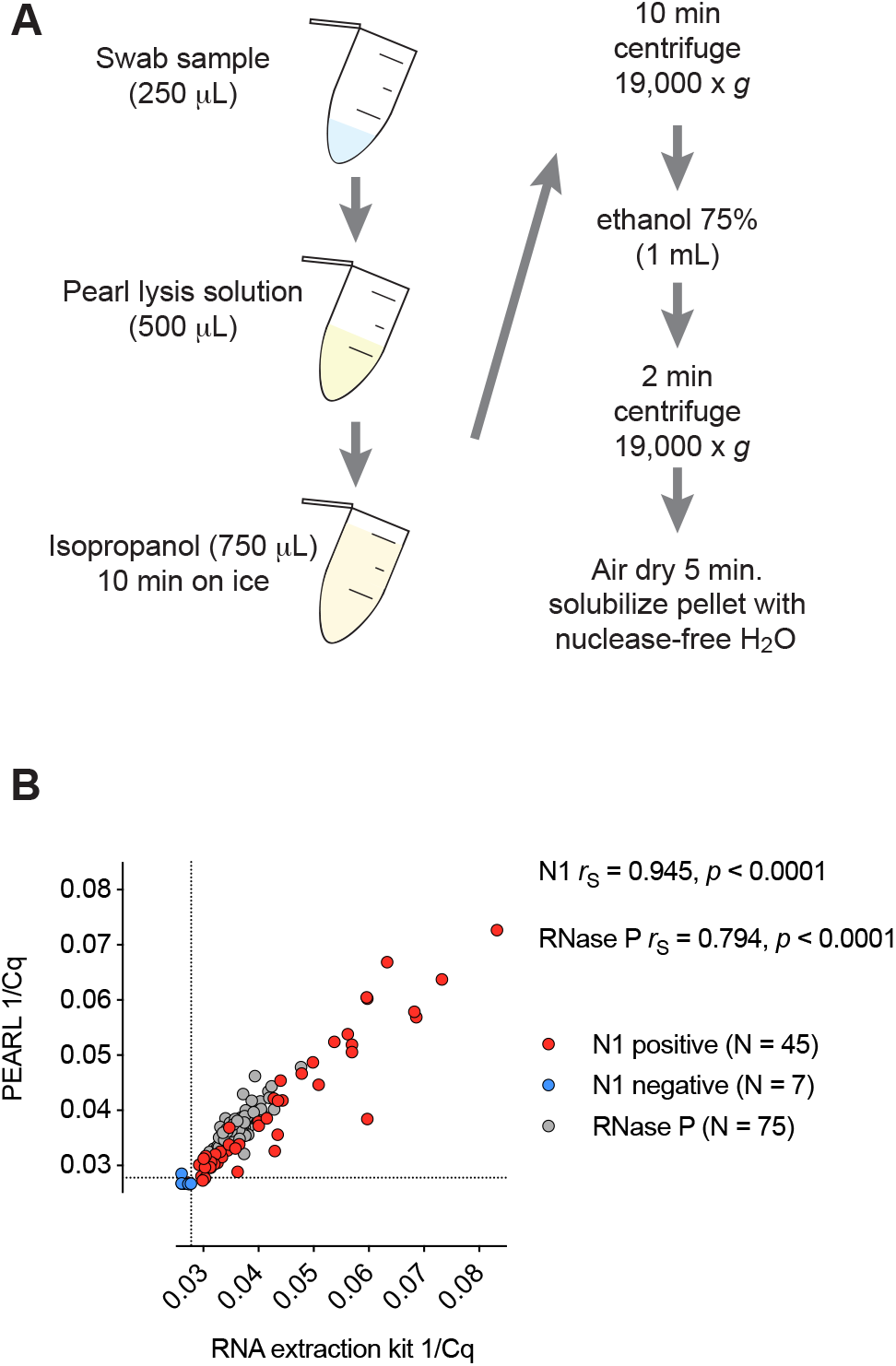
**A)** Overview of PEARL. **B)** Comparative RT-qPCR analysis of the levels of SARS-CoV-2 nucleocapsid (N1) and RNaseP RNA sequences in de-identified SARS-CoV-2 positive and negative samples after RNA extraction using PEARL or an RNA extraction kit (QIAamp Mini Elute Virus Spin Kit) in nasopharyngeal swab samples. Dotted lines indicate a limit of detection of 36 cycles. We obtained Cq values for N1 below our limit of detection in 7 negative samples out of 30. No Cq values were obtained for N1 in the remaining 23 samples.

To maximize SARS-CoV-2 detection sensitivity, we tested various ratios of sample to PEARL lysis solution. In these experiments, we observed that 250 μL of initial swab sample input and 500 μL of PEARL lysis solution resulted in the lowest RT-qPCR Cq values (Fig. S1). Next, we compared the Cq values from PEARL extracts to RNA purified using a commercial kit. We found strong concordance between extraction methods using de-identified nasopharyngeal swab samples provided by the Santa Barbara County Public Health Department (Fig. 1B). However, PEARL, required a modest increase in initial sample input (1.25-fold) to achieve similar sensitivity to that of the commercial RNA extraction kit we used (Fig. 1B, note that the sample input for PEARL was either 175 μL or 250 μL, while the sample input for the commercial kit we used was 200 μL or 140 μL, respectively). Together, these results indicate that PEARL can be used as an alternative to commercial RNA extraction kits without substantial loss in sensitivity.

We reasoned that, because DNA and protein co-precipitate with RNA upon addition of isopropanol during extraction^12^, PEARL could be used to streamline the retrieval of RNA, DNA, and proteins from other viruses. To test whether PEARL can be used to detect different types of viruses, we used cells infected with the Kaposi’s Sarcoma Associated Herpesvirus (KSHV), which contains a DNA genome, or with Zika virus (ZIKV), a flavivirus that contains an RNA genome and no DNA replication intermediates in its life cycle^13^. In these experiments, we used iSLK-219 cells, which are latently infected with a GFP-expressing recombinant KSHV^14^, or HeLa cells infected with the PRVABC59 strain of ZIKV, which was isolated in Puerto Rico in 2015^15^, at a multiplicity of infection (MOI) of 1. We collected 100,000 cells, which corresponds to the estimated cellular yield of a typical buccal swab^16^, and prepared 10-fold dilutions to determine the detection limit for RNA, DNA, and protein. Next, we prepared PEARL extracts and probed for viral and host nucleic acids and proteins using qPCR- and immunodetection-based assays, respectively. To ensure the specificity of RNA or DNA detection, we treated the PEARL extracts with DNase I (to detect RNA) or RNase A (to detect DNA). For protein immunodetection, we treated the PEARL extracts with RNase and DNase before SDS-PAGE and western blotting to ensure undisturbed migration of the proteins during electrophoresis, or left them untreated for dot-blot detection.

To detect host and viral transcripts, we synthesized first-strand complementary DNA from the DNase I treated samples and used it for qPCR detection of the host β-actin mRNA (*ACTB*) and viral transcripts. These viral mRNAs included the KSHV latency-associated nuclear antigen (*LANA*) and KHSV-encoded *GFP* (Fig. 2A), as well as ZIKV RNA regions encoding the non-structural proteins NS1 and NS5 (Fig. 3A). In these experiments, we detected viral transcripts in PEARL extracts obtained from as few as 1,000 infected cells, and we did not observe significant differences in sensitivity between the detection of KSHV or ZIKV transcripts. Thus, PEARL can facilitate the detection of mRNAs from DNA and RNA viruses.

**Figure 2.**
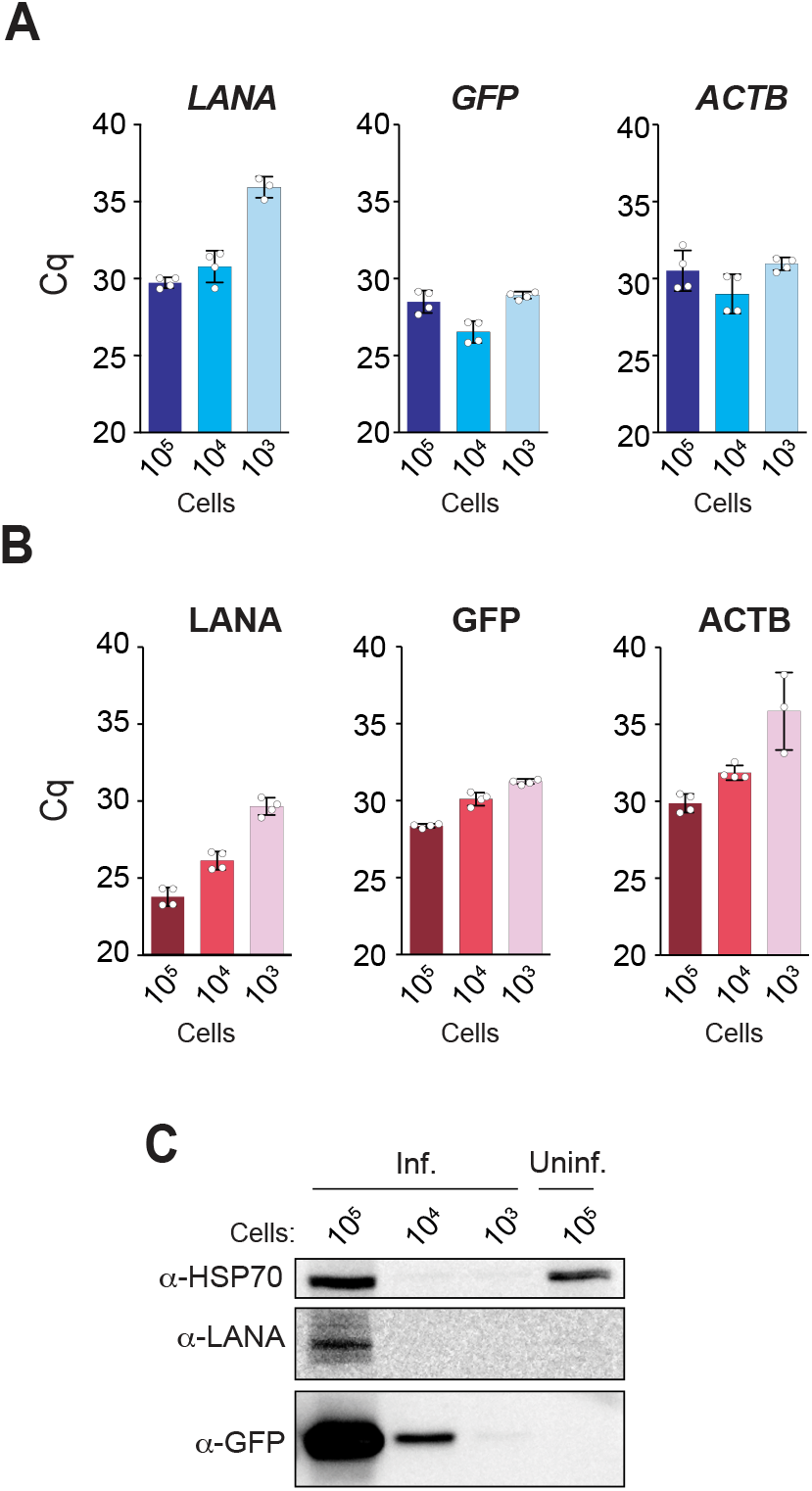
**A)** RT-qPCR analysis of the levels of KSHV (LANA, GFP) and host β-actin (ACTB) mRNAs, and **B)** their corresponding genomic sequences. **C)** Western blot analysis of the expression of KSHV (LANA, GFP) and host (HSP70) proteins. Inf., infected; Uninf., uninfected.

**Figure 3.**
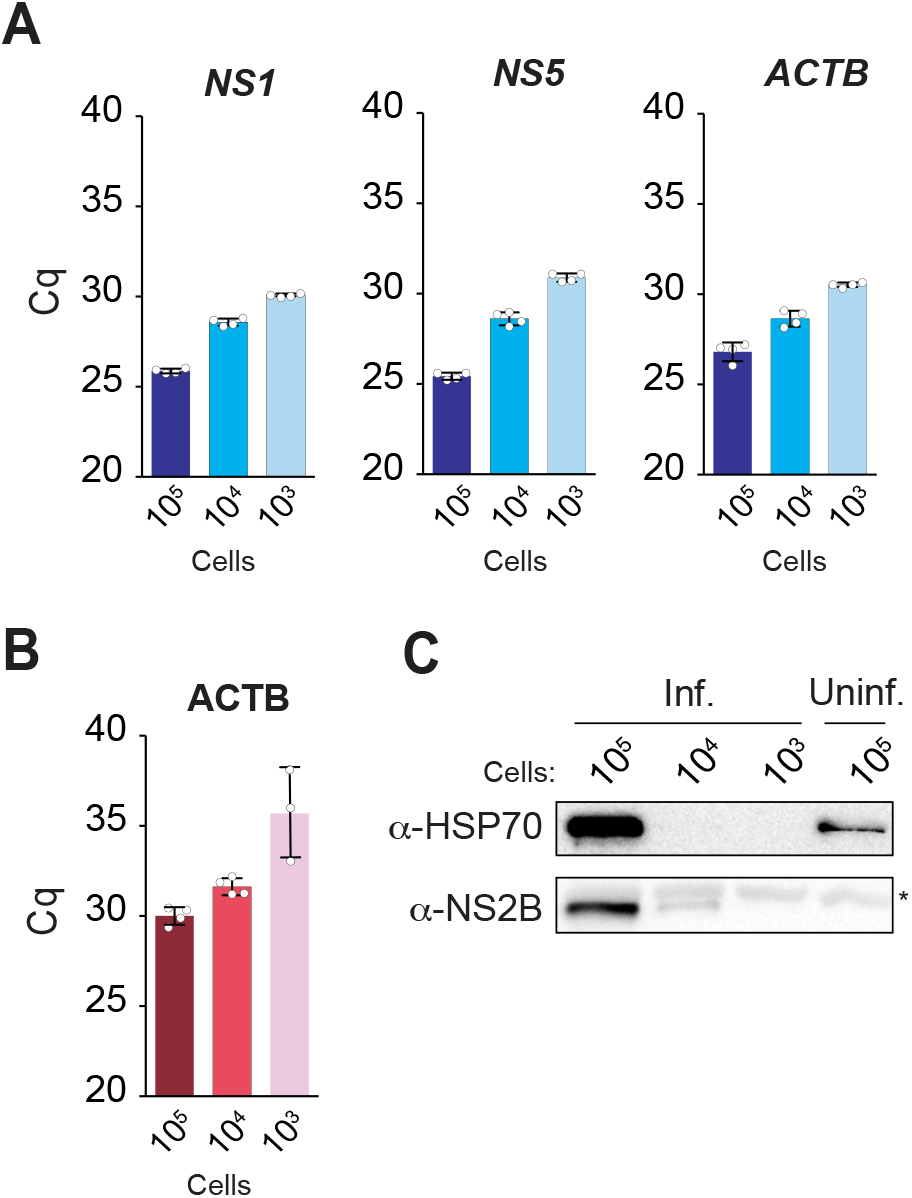
**A)** RT-qPCR analysis of the levels of ZIKV non-structural proteins NS1 and NS5 and host β-actin (ACTB) mRNAs **B)** qPCR analysis of the expression host ACTB genomic DNA sequences in ZIKV-infected samples. **C)** Western blot analysis of expression of ZIKV (NS2B) and host (HSP70) proteins. *non-specific band.

To detect host and KSHV genomic sequences, we used PEARL extracts treated with RNase A. Our target sequences for DNA detection corresponded to the genes for the host and viral transcripts aforementioned (Fig. 2A, 3A). In agreement with our observations for KSHV transcripts, we detected the viral genome in as low as 1,000 latently infected cells (Fig 2B). We also detected the host DNA β-Actin locus in all samples, regardless of the infection status (Fig 2B, 3B). In these experiments, we used the same pair of PCR primers for detection of the β-Actin mRNA and genomic DNA sequences, thus eliminating variability that could arise from dissimilar amplification efficiencies of different primer pairs. The primers target sequences in different β-Actin exons (Fig. S2A), distinguishing mRNA amplicons from genomic DNA amplicons by molecular size. As a control, and to corroborate that the amplification products in Figs. 2B and 3B were not templated on contaminant RNA, we used PCR primers that amplify the non-transcribed promoter region of the host gene HSPA5 (Fig. S2B). As expected, we detected an amplification product only in the PEARL extracts treated with RNase A but not in those treated with DNase I (Figs. S2C, S2D), verifying the specificity of the amplification reaction.

An additional benefit of PEARL over column-based commercially available RNA extraction methods is that it allows the recovery of proteins in addition to nucleic acids. To confirm the presence of host and viral proteins in PEARL extracts, we carried out western blot and dot-blot assays using antibodies against the ubiquitous host chaperone HSP70, and the viral proteins KHSV-LANA, KSHV-encoded GFP, and ZIKV-NS2B. In these experiments, we detected the host chaperone HSP70 in PEARL extracts obtained from 100,000 cells (HeLa and iSLK-219) by western blot (Fig. 2C and 3C) and in as few as 12,500 cells (HeLa and iSLK-219) by dot-blot (Fig. S3A, and S3B). Detection of KSHV-encoded GFP was achievable with approximately 1,000 iSLK-219 cells (Figs. 2C and S3B). Detection of KSHV-LANA was significantly less sensitive by western blot than by dot-blot, requiring 100,000 and 1,250 iSLK-219 cells, respectively (Figs. 2C and S3B). Taken together, our results indicate that PEARL can be used as a reliable and efficient method to extract host and virus nucleic acids and proteins from a wide range of viral infections.

While we designed PEARL to be accessible, it still uses a high-speed centrifuge, which is expensive, requires AC power to operate, and is typically restricted to professional laboratories. We reasoned that we could make PEARL field deployable by using a hand-powered centrifugation device. Inspired by the work of Manu Prakash and others who pioneered these types of devices^17–19^, we modified a freely-available design for a hand-powered centrifuge actuated by supercoiling of a string (https://www.thingiverse.com/thing:1946291). We engineered a safety lid and chamfered all edges to avoid abrasion on the string, and increased the distance between the pull points to augment torque around the rotational axis (Fig. S4A). We 3D printed our device with thermoplastic polyester, measured its angular velocity using a laser tachometer, and found that we could achieve centrifugal forces of approximately 3,900 × *g* (Fig. S4B). To test whether our hand-powered centrifuge could replace a benchtop centrifuge in PEARL, we compared the RNA and protein extraction efficiency achieved with our device and with a tabletop centrifuge set at 19,000 × *g.* Despite the substantial difference in centrifugal force, we found that viral RNA and proteins can be easily detected using PEARL extracts prepared with our hand-powered centrifuge (Figs. S4C and S4D). Thus, this device can enable the deployment of PEARL in the field without a significant loss in detection sensitivity.

## Discussion

The primary tool to combat the COVID-19 pandemic is widespread and accessible testing to monitor SARS-CoV-2 prevalence and spread, which informs deployment of containment and mitigation measures^20^. Globally scaled testing remains an unmet public health need, as attempts to meet this demand have resulted in shortages of the reagents and supplies necessary for sample processing, RNA extraction and SARS-CoV-2 detection. Here we present data to support PEARL as a cost effective, simple, and less-toxic alternative for the isolation of RNA, DNA and proteins. Our results indicate that PEARL enables the detection of SARS-CoV-2 transcripts in COVID-19 positive swab samples with sensitivity comparable to that afforded by commercially available RNA extraction kits. This outcome highlights the validity of using PEARL as a viable alternative to facilitate the detection of SARS-CoV-2 in respiratory samples.

Our data also show that PEARL extracts can be used to efficiently detect host and viral transcripts, genomic DNA, and proteins regardless of the nature of the infection—PEARL was equally useful in detecting DNA and RNA viruses with different tropism. Coupling PEARL to different downstream analyses for detection of nucleic acids and proteins can provide a powerful tool for detection of diverse viruses. Moreover, because RNA, DNA, and proteins are extracted at once, PEARL reduces sample handling time, allowing for streamlined diagnostic procedures. Thus, it may enable both nucleic acid and antigen-based SARS-CoV-2 testing. PEARL’s minimal handling requirements also make it scalable, which is desirable for high-volume testing operations, as is needed for SARS-CoV-2 testing.

It is possible that the collection medium used to store samples before processing may influence the performance of PEARL. For example, the viral transport media recommended by the CDC to store and inactivate samples for SARS-CoV-2 testing (2% Fetal Bovine Serum, 100 μg/mL Gentamicin, 0.5 μg/mL Amphotericin B, and various salts)^21^ has components that could co-precipitate with target analytes. Isopropanol is less polar than ethanol, and therefore, it has a higher propensity to precipitate salts and antibiotics^22^. In our experiments, co-precipitation of salts and antibiotics does not appear to compromise downstream RT-qPCR or immunodetection assays. Concerns regarding the inhibition of downstream detection assays could be addressed by using ethanol instead of isopropanol.

It is also possible that PEARL may introduce extraction bias, as short RNAs, including tRNAs, snoRNAs and miRNAs, are more difficult to precipitate than longer RNA and DNA molecules^22^. Though we have not directly tested whether small RNAs are underrepresented in PEARL extracts, we have designed PEARL to enhance the precipitation of all RNAs by using linear polyacrylamide as a carrier^23^. Additionally, longer and faster centrifugation speeds can be used to enhance small RNA recovery, if needed10. Further improvements may be required to implement PEARL as mainstream nucleic acid and protein isolation tool for detection of viruses obtained from sources different than those described here, as the sample type may dictate overall performance. Future work outside the scope of this study will be required to address whether this is the case.

Finally, since PEARL uses common reagents and it does not require expensive equipment or highly trained personnel, it can provide an accessible alternative for streamlining diagnostics in geographic areas that lack access to capital, specialized reagents, and professional laboratories. Moreover, PEARL is field-deployable, given that a hand-powered centrifugation device can be used. In view of these considerations, coupling PEARL to our recently developed CRISPR-based protocol for detection of SARS-CoV-2 genetic material called CREST (Cas13-based, Rugged, Equitable, Scalable Testing)^24^ could allow efficient, affordable, widespread testing, lowering the barrier of “luxury testing” in many regions of the world.

## Materials and Methods

### PEARL

Samples were mixed in a 1:2 (v/v) sample:lysis solution (0.5% IGEPAL CA-630, 450 mM sodium acetate, 20 % glycerol, 20 mM TCEP, 50 μg/ml linear polyacrylamide, and 20 mM HEPES-KOH pH 7.2) ratio and incubated for 5-minutes at room temperature. Next, nucleic acids and proteins were precipitated on ice, for 10 minutes, using one volume of cold isopropanol. The precipitated material was collected by centrifugation at 19,000 × *g* for 10 minutes, washed once with 75% ethanol, air-dried for 5 minutes at room temperature and solubilized in 20 μl of nuclease-free water for amplification-based detection of nucleic acids or immunodetection of proteins.

### Cell culture and infections

All cell lines were maintained in Dulbecco’s modified Eagle’s medium (DMEM) supplemented with 10% of fetal bovine serum (FBS), L-Glutamine, and antibiotics (penicillin/streptomycin, 100 units/mL), and were maintained in a humidified incubator at 37°C and 5% CO_2_. iSLK-219 cells are latently infected with KSHV.21914. This recombinant virus is maintained in cells as an episome. GFP is constitutively expressed from the episome, under the control of the human EF1 promoter. iSLK-219 cells also harbor the gene for a doxycycline-inducible KSHV RTA transcription activator. Uninfected iSLK and KSHV-infected iSLK-219 cells were grown to 80% confluence, collected by trypsinization after two washes with PBS (GenClone), counted, and diluted at the desired density in 250 μL of PBS for PEARL extraction. For ZIKV infections, HeLa cells were grown to 60% confluency and then infected with ZIKV at a multiplicity of infection (MOI) of 1. 48 hours post-infection, the cells were collected by trypsinization after two washes with PBS (GenClone), and counted. Cells were diluted in 250 μL of PBS for PEARL extraction.

### qPCR

PEARL extracts were obtained from de-identified human samples or cultured cells. SARS-CoV-2 positive human samples were heat-inactivated by incubation at 56 °C for 30 minutes before RNA extraction. RNA from these samples was obtained with the QIAamp Mini Elute Virus Spin Kit (Qiagen) following the manufacturer’s protocol, using 200 μL of sample input and eluting the purified RNA in 50 μL. PEARL extracts were prepared using 250 μL of SARS-CoV-2 positive human samples or a fixed number of cultured infected cells suspended in 250 μL of PBS. PEARL extracts from cultured cells were treated with either DNase I (1 unit per every 8 μL of PEARL extract, New England BioLabs) or with RNase A (0.1 mg per every 8 μL of PEARL extract, ThermoFisher) in a final volume of 10 μL for 30 minutes at 37 °C. 5 μL of DNase-treated samples were reverse transcribed in a final volume of 10 μL using the iScript cDNA synthesis kit (Bio-Rad) following the manufacturer’s protocol and diluted 5-fold in nuclease-free water before qPCR. Target detection by qPCR was carried out with SYBR Select Master Mix (Applied Biosystems) using 2 μL of diluted cDNA as template, and following the manufacturer’s protocol. The entire 10 μL from RNase-treated samples (genomic DNA) were diluted 5-fold with nuclease-free water and 2 μL of diluted genomic DNA were used as template for detection of specific genes with the SYBR Select Master Mix (Applied Biosystems) following the manufacturer’s protocol. Detection of SARS-CoV-2 N1 gene sequences and host RNase P mRNA from de-identified SARS-CoV-2 positive samples was carried out with the one-step TaqMan RNA-to-Ct 1-Step Kit (ThermoFisher), using 2 μL of undiluted PEARL extract, and following the manufacturer’s protocol. All qPCR data were collected using a CFX96 touch real-time PCR instrument (BioRad), and analyzed with the CFX Maestro 1.1 software (BioRad). Cq values were determined by regression. Data analysis and statistical tests were performed using the Graph Pad Prism 6.0 software.

### Immunodetection

Nuclease-treated PEARL extracts were separated on 10% SDS-PAGE gels and transferred onto nitrocellulose membranes (Bio-Rad) for western blot analysis. The membranes were blocked in 0.5% BSA-TBST for 30 minutes. Primary antibodies were diluted in 0.5% BSA-TBST as follows: anti-HSP-70 (Cell Signaling Technology 4872) 1:1,000; anti-LANA/ORF73 (Advanced Biotechnologies 13-210-100) 1:3,000; anti-GFP (Invitrogen A11122) 1:3,000; anti-NS2B (GeneTex GTX133308) 1:1,000. The membranes were incubated with primary antibodies for 30 minutes at room temperature. Following primary antibody incubation, the membranes were washed with TBST 3 times before the addition of HRP-conjugated secondary antibodies. The membranes were incubated for 30 minutes with secondary antibody diluted 1:3,000 in 0.5% BSA-TBST. Immunoreactivity was detected using the Radiance Plus HRP Substrate (Azure Biosystems). All images were captured with an Azure Biosystems C300 gel imaging system. Image post-processing was carried out in Photoshop CC (Adobe) using automatic contrast. For dot-blot-based immunodetection, nitrocellulose membranes (Bio-Rad) were spotted with 1 μL of PEARL extract and allowed to dry completely at room temperature for 30 minutes. For the remainder of the procedure, membranes were processed and imaged as described for western blotting.

### Hand-powered centrifuge

Our hand-powered centrifuge was designed in SolidWorks 2018 (Dassault Systemes), sliced (0.2 mm layer height) in Cura (Ultimaker), and printed on an Ender3 3D printer (Creality) using 1.75 mm polylactic acid filament (Hatchbox Inc). To actuate our device, we used Brutal Strong 135-test braided fishing line (Izorline International). Approximately 1 m of line was threaded through holes designed for the string-driven system in the hand pulls and in the rotor, and the line was secured to itself with a double uni-knot forming a loop. Maximum angular speed was determined by affixing reflective tape to the rotor and revolutions per minute (RPM) were measured with a laser tachometer (Neiko) over a 1 second sampling time. The maximum relative centrifugal force (RCF) was calculated as follows:

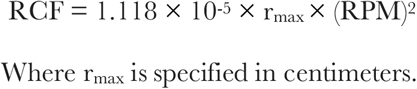

3D print files can be found at: https://3dprint.nih.gov/discover/3dpx-014683. After PEARL precipitation, the samples were spun at maximum speed with the hand-powered centrifuge, or at 19,000 × *g* in a benchtop centrifuge. RNA and protein recovery for both centrifugation methods were determined by RT-qPCR and dot blot.

## Acknowledgements

We would like to thank Jennifer Rauch, Sabrina Solley, Ryan Lach, and Eric Valois for insightful discussions. We thank Stewart W. Comer of the Santa Barbara County Public Health Department for granting us access to SARS-CoV-2 positive samples. We thank Markus Merk for donating PLA filament for 3D printing, and David Asplund and Olivia Quinn for their assistance with RPM determination of the hand-powered centrifuge. We thank the UCSB Office of Research for their generous support. We are thankful to our colleagues, our University directives, and all the health workers, without whom this work would have not been possible.

