## Supplemental Tables and Figures for "A Fast and Accessible Method for the Isolation of RNA, DNA, and Protein to Facilitate the Detection of SARS-CoV-2"

---

| Reagent | Cost (USD) | USD/gram/mL | For 500μL | Cost/reaction |
|---|---|---|---|---|
| HEPES-KOH | $728/1 Kg | $0.73 | 0.002383 g | $0.00173482 |
| NaOAc | $329.5/1 Kg | $0.33 | 0.0150075 g | $0.00494497 |
| Igepal CA-630 | $98.2/500 mL | $0.1964 | 0.0025 mL | $0.000491 |
| Glycerol | $113/500 mL | $0.22675 | 0.2 mL | $0.04535 |
| TCEP | $178/10 g | $17.8 | 0.0028665 g | $0.0510237 |
| Polyacrylamide | $173/250 g | $0.692 | 0.000025 g | $0.0000173 |
| Isopropanol | $50.15/500 mL | $0.1003 | 0.75 mL | $0.075 |
| Total | | | | $0.17851 |

**Supplemental Table 1.** Cost breakdown of PEARL reagents. Note that the cost of one PEARL extraction is approximately 20 times lower than column-based RNA extraction.

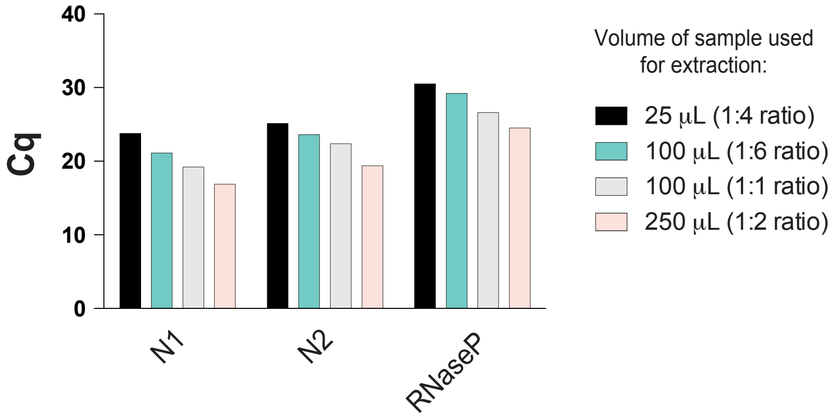


**Supplemental Figure S1.** Threshold values obtained by RT-qPCR for SARS-CoV-2 N1 and N2 viral transcripts, and the RNase P host mRNA, obtained using PEARL extracts. Different volumes of swab samples with the indicated sample to PEARL-lysis-buffer ratios were used to determine the conditions that maximize readout.

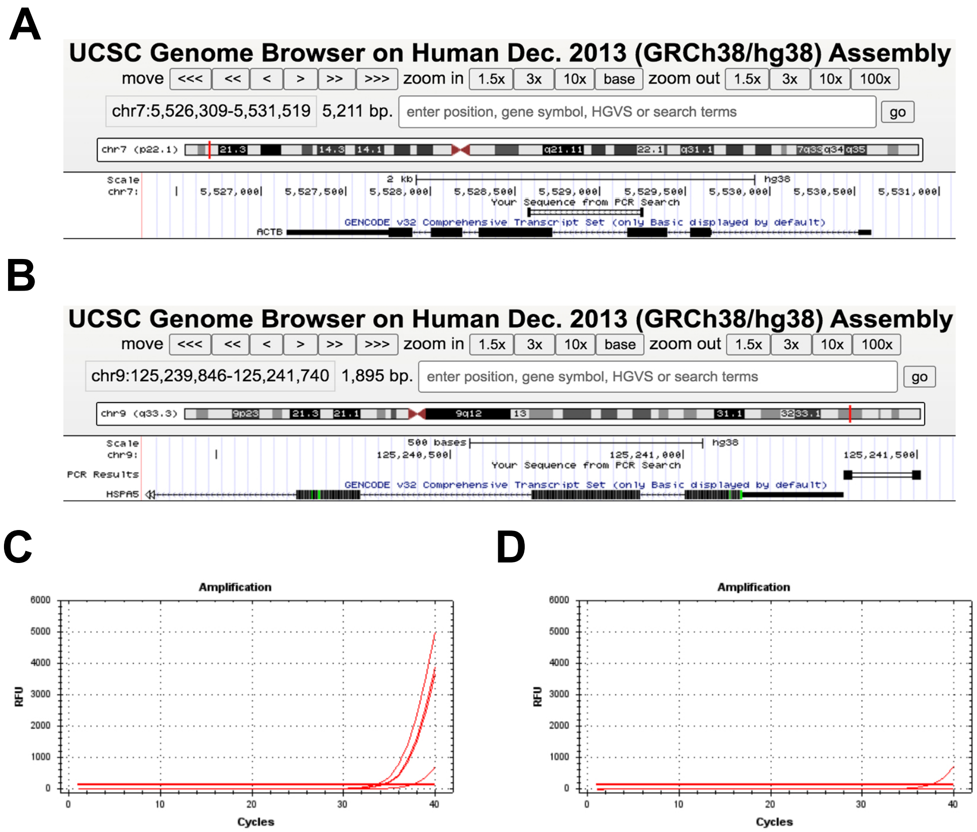


**Supplemental Figure S2.** Genomic locations of A) PCR products obtained with oligonucleotides targeting exons 3 and 4 in the ACTB locus and B) a PCR product corresponding to the HSPA5 proximal promoter region. Tracks were obtained by *in silico* PCR and were mapped onto the latest assembly of the human genome in the UCSC genome browser (https://genome.ucsc.edu/index.html). C) and D) Amplification curves for the HSPA5 promoter region using the PCR primers shown in B in using PEARL extracts treated with (C) RNase A or (D) DNase I.

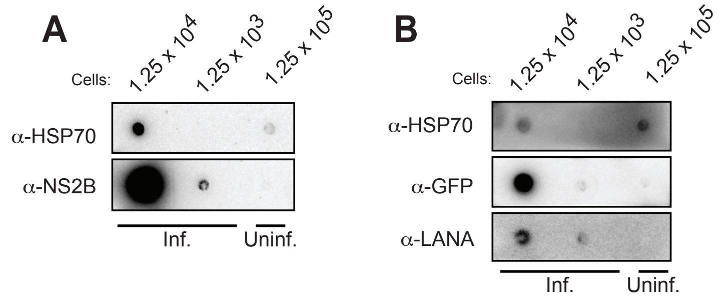


**Supplemental Figure S3**. A) Dot blot immunodetection of ZIKV (NS2B) and host (HSP70) proteins. B) Dot blot immunodetection of KSHV (GFP, LANA) and host (HSP70) proteins. Inf., infected; Uninf., uninfected.

A

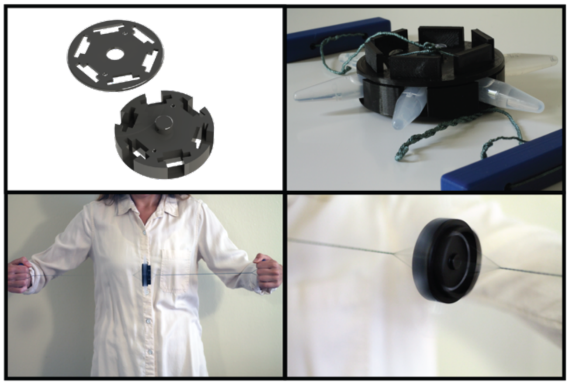

B

| Pull # | RPM (Max) | RCF (Max) |
|---|---|---|
| 1 | 7240 | 3809 |
| 2 | 7324 | 3898 |
| 3 | 6915 | 3475 |
| 4 | 7812 | 4435 |
| 5 | 7547 | 4139 |
| 6 | 6432 | 3006 |
| 7 | 6665 | 3228 |
| 8 | 7836 | 4462 |
| 9 | 6846 | 3406 |
| 10 | 8534 | 5292 |
| **Average** | **7315** | **3915** |

C

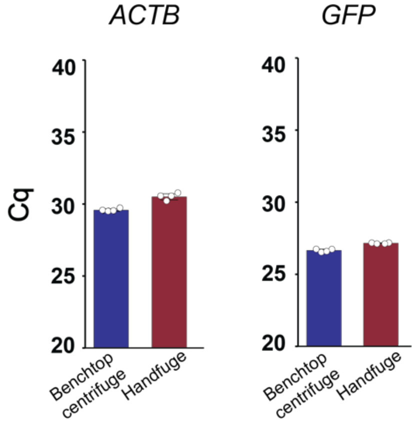


D

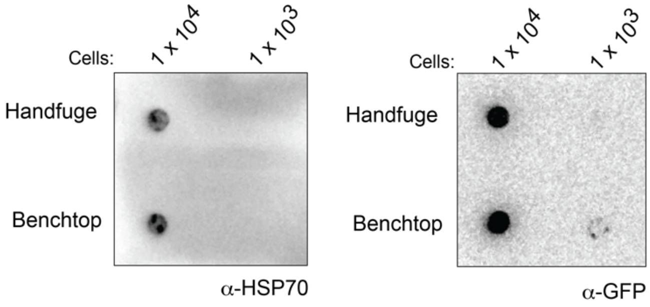


**Supplemental Figure S4**. A) Top left: 3D render of our hand-powered centrifuge. CAD files can be found here: https://3dprint.nih.gov/discover/3dpx-014683. Top right: Assembled hand-powered centrifuge with its rotor loaded and the driving strings supercoiled. Bottom left: Actuation of the hand-powered centrifuge by string supercoiling. Bottom right: Close-up image of the spinning rotor during actuation. B) Laser tachometer data averaged from 10 actuation events of the hand-powered centrifuge. C) Cq values obtained by probing for host and KSHV mRNAs using RT-qPCR using PEARL extracts prepared with either a benchtop laboratory centrifuge or our hand-powered centrifuge, “Handfuge”. D) Dot blots obtained by probing for host and KSHV proteins in PEARL extracts prepared as in C.

*List of oligonucleotides used for qPCR in this study*

| Target | Organism | Forward oligonucleotide | Reverse oligonucleotide |
|---|---|---|---|
| ACTB | *Homo Sapiens* | TTCTACAATGAGCTGCGTGTG | AGGGCATACCCCTCGTAGAT |
| HSP5A (promoter) | *Homo Sapiens* | GCGGAGCAGTGACGTTTATT | ACCTCACCGTCGCCTACTC |
| NS1 | Zika virus PRABC59 | ATAACAGCTTTGTCGTGGATG | TAACCTTGAGCCAGACACTAG |
| NS5 | Zika virus PRABC59 | GACTGGGTTCCAACTGGGAG | CCACACTCTGTTCCACACCA |
| GFP | KSHV | GAGCGCACCATCTTCTTCAAG | GGCGGATCTTGAAGTTCAC |
| LANA | KSHV | CCTCCATCCCATCCTGTGTC | GGACGCATAGGTGTTGAAGAG |
